# Microstructural Spine Alterations Increase Neuronal Excitability in Focal Cortical Dysplasia

**DOI:** 10.1101/2025.10.16.682987

**Authors:** Jawon Gim, Na-Young Seo, Gyu Hyun Kim, Kea Joo Lee, Joon Ho Choi

## Abstract

Focal cortical dysplasia(FCD) is a leading cause of drug-resistant epilepsy, traditionally attributed to impaired inhibition. However, recent ultrastructural evidence suggests excitatory synaptic alterations may also contribute. To investigate this, we performed computational modeling of human pyramidal neurons in NEURON, incorporating electron microscopy–derived dendritic spine morphologies. Baseline and FCD-like models were constructed with differences in spine density and morphology while maintaining comparable net synaptic input.

Simulations revealed that reduced spine density increased input resistance and amplified excitatory postsynaptic potentials, enhancing dendritic-to-somatic transmission. Spine neck geometry, but not head size, critically shaped voltage compartmentalization. The altered model required markedly fewer synchronous inputs to reach spiking threshold and exhibited higher firing rates under Poisson-distributed synaptic activity, particularly under sparse input conditions.

These results demonstrate that excitatory microstructural changes - reduced spine density and weakened neck compartmentalization - can elevate neuronal excitability, in addition to the inhibitory deficit which is the biggest known effect. These findings highlight excitatory spine remodeling as another key driver of epileptogenesis. We speculate that extending this analysis will be essential to fully understand circuit-level hyperexcitability and to identify new therapeutic targets.

## Introduction

Focal cortical dysplasia (FCD) is a severe malformation of neocortical development, and is one of the leading causes of drug-resistant epilepsy. Due to its pharmaco-resistant nature, surgical intervention is often required for effective treatment^1-3^. FCD is characterized by abnormal cortical lamination and disorganized neuronal architecture, often accompanied by cytological abnormalities such as dysmorphic neurons and balloon cells. Such disruptions alter cortical circuit dynamics and establish FCD as a significant substrate for epileptogenesis.^4,5^. Electrophysiological recordings, immunohistochemical staining and computational modeling studies consistently show disruption of the excitation-inhibition(E-I) balance within tissue from a patient with FCD ^6^. Notably, a reduction in inhibitory signaling has been considered the main factor of epileptogenesis^6-12^.

However, this finding does not fully capture the complexity of the epileptogenic mechanism. Some computational studies suggest that epileptic activity can be generated without such reduced inhibition^13,14^. Despite differing views on the role of inhibitory mechanisms, studies converge on a well-known key scenario: a scheme that input to output firing rate relation is increased in FCD affected area compared to normal cortex.

Adding further complexity, a recent high-resolution electron microscopy(EM) study revealed that not only inhibitory synapses but also excitatory synapses show reduced density on pyramidal neuron^15^. An interesting feature of that was the existence of extra-large spines and its excitatory synapses which contain a huge number of synaptic vesicles. These features give rise to the counterintuitive question of whether the decreased number of synapses also lowers firing rate of pyramidal neurons or the increased size of each synapse raises the firing rate of that neuron.

Because the decrease of inhibitory synapse density raises firing rate for sure, to test the outcome of alteration in excitatory synapse are not easy to isolate experimentally. Thus we performed a preliminary mathematical analysis of such neuron types with altered synaptic density and size while preserving the same net input current. It showed a higher firing rate in epileptic neurons compared to control ones. To validate this, we designed computational model experiments with morphological details of a human pyramidal neuron using NEURON^16^. On that model cell, the effect of spine density and size alterations observed in samples of a patient with FCD are tested.

Taken together, our findings underscore the importance of both synaptic number and strength in shaping cortical excitability in FCD. These findings suggest that epileptogenesis arises from the convergence of inhibitory deficits and excitatory synaptic spine remodeling. Understanding how these alterations jointly shape network activity may provide new mechanistic insights into the cortical hyperexcitability mechanism and potential therapeutic strategies.

## Methods

### Simulation

All simulations were conducted in NEURON (v8.2)^16^ with Python (v3.10)^17^.

### Model cell construction

Baseline and altered model cells were constructed from morphological features of human pyramidal neurons^18^ (Figure 1A). Briefly, each cell included a spherical soma (17μm diameter), six basal dendrites (1μm × 381μm), and a single apical trunk (30μm long) tapering into a tuft of 14 apical dendrites (1μm × 351μm). Dendritic spines were uniformly distributed beyond 30μm from the soma, with densities derived from volume EM data^15^ (Figure 1B,C). On the most distal 101μm of each dendrite, spines with realistic neck and head dimensions were explicitly modeled and randomly positioned (Figure 1D). All dendritic and spine compartments were passive; the soma contained Hodgkin–Huxley-type Na+ and K+ channels^19^. Axial and membrane resistances, capacitance, and channel conductances followed tested values^19-21^. Alpha-amino-3-hydroxy-5-methyl-4-isoxazolepropionic acid (AMPA) and N-methyl-D-aspartate (NMDA) receptors were assigned to explicit spines using conductance parameters reported in various studies^7,22^.

**Figure 1.**
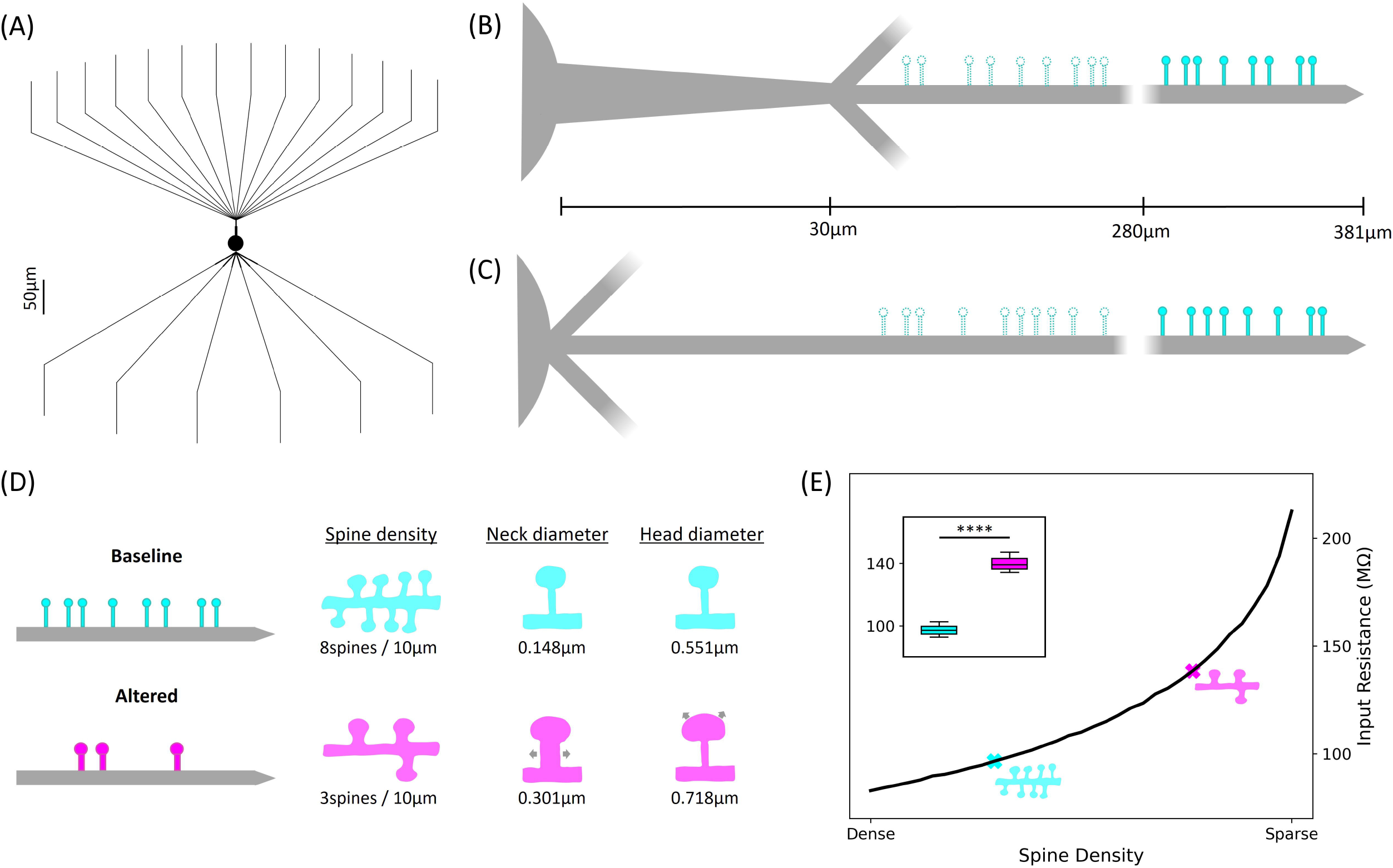
Dendritic morphology and input resistance across models. (A) Whole-cell morphology of the baseline and altered models. Apical (B) and basal (C) dendrites showing distribution of implicitly placed (dashed) and explicitly placed (solid) spines. A portion of each dendrite is omitted to emphasize differences between models. Horizontal lines indicate distance from the soma. (D) Distal dendritic spine morphologies in the two models: baseline in cyan, altered in magenta. (E) Input resistance as a function of spine density, with model-specific positions indicated (cyan: baseline; magenta: altered). Inset box plot shows input resistance across 10 configurations per model. **** P < 0.0001, unpaired two-tailed t-test.

### Measurement of input resistance

Input resistance was determined by injecting a 0.1nA, 500ms somatic current and dividing the resulting steady-state voltage by the current.

### Modeling of spine density and morphological scaling

Changes in dendritic surface area from spines were incorporated using a spine factor^23^. Based on reconstructed data^15^, spine factors of 2.0 (baseline) and 1.2 (altered) were applied to all dendrites except the explicitly modeled distal segments.

### Single-Spine EPSP Simulations

To assess spine morphology effects, a single dendritic spine was randomly placed and stimulated via AMPA and NMDA receptors with fixed conductance. Membrane potentials were recorded at the spine head, spine base, and soma (Supplementary Figure 1A). In simulations of neck diameter, head diameter, or spine density, stimulated spine locations ranged from proximal to distal dendritic segments to capture morphology-dependent effects. Each condition was simulated 100 times to obtain EPSP distributions.

### Spiking Threshold via Synchronous Activation

In each trial, the explicit spines distributed along the distal dendrite were randomized. From this randomized set, a specified number of spines (N) was randomly selected and activated simultaneously. For each value of N, 20 independent simulations were performed.

### Poisson-Distributed Synaptic Input

Excitability was tested using independent Poisson spike trains (0.2–5Hz) assigned to each spine to mimic asynchronous input. Synaptic conductances were set based on prior electrophysiological data ^7^. Each simulation lasted 12s, and spikes between 1–10s were counted to compute firing rates. Ten distinct apical/basal dendrite configurations were generated, with three independent runs per configuration per rate, totaling 30 simulations per rate with randomized spine assignments in each trial.

### Code availability

code available from https://github.com/jawonGim/Simulation_SpineAlterationFocalCorticalDysplasia.git

## Results

### Validation of model cell properties

The baseline properties of the model cells were verified by measuring their input resistance (Figure 1E; see Methods). The resulting values were comparable to those reported for human layer 2/3 pyramidal neurons^7^.

### Impact of Spine Density and Morphology on EPSP Propagation and Neuronal Excitability

To investigate how dendritic spine architecture affects neuronal excitability, we simulated the impact of spine density, neck diameter, and head diameter on single-spine–evoked excitatory postsynaptic potentials (EPSPs).

The altered model, with a 2.6-fold lower spine density, exhibited consistently larger EPSP amplitudes across all recording sites (Figure 2A, Supplementary Figure 1B,C), indicating that reduced spine density enhances synaptic potentiation. A continuum of model neurons spanning a range of spine densities revealed that input resistance increased monotonically as spine density decreased (Figure 1E), suggesting that lower spine density elevates global excitability by amplifying responses to synaptic input. While EPSP differences at the spine head and base were mainly magnitude-dependent, somatic EPSPs in the altered model showed a broader upward shift (Figure 2A). Analysis of somatic EPSP amplitude relative to the distance from soma to the stimulated spine (Figure 2B) showed that signal attenuation was 62% in the baseline model versus 44% in the altered model, reflecting more increased dendritic-to-somatic signal transmission and elevated excitability with reduced spine density.

**Figure 2.**
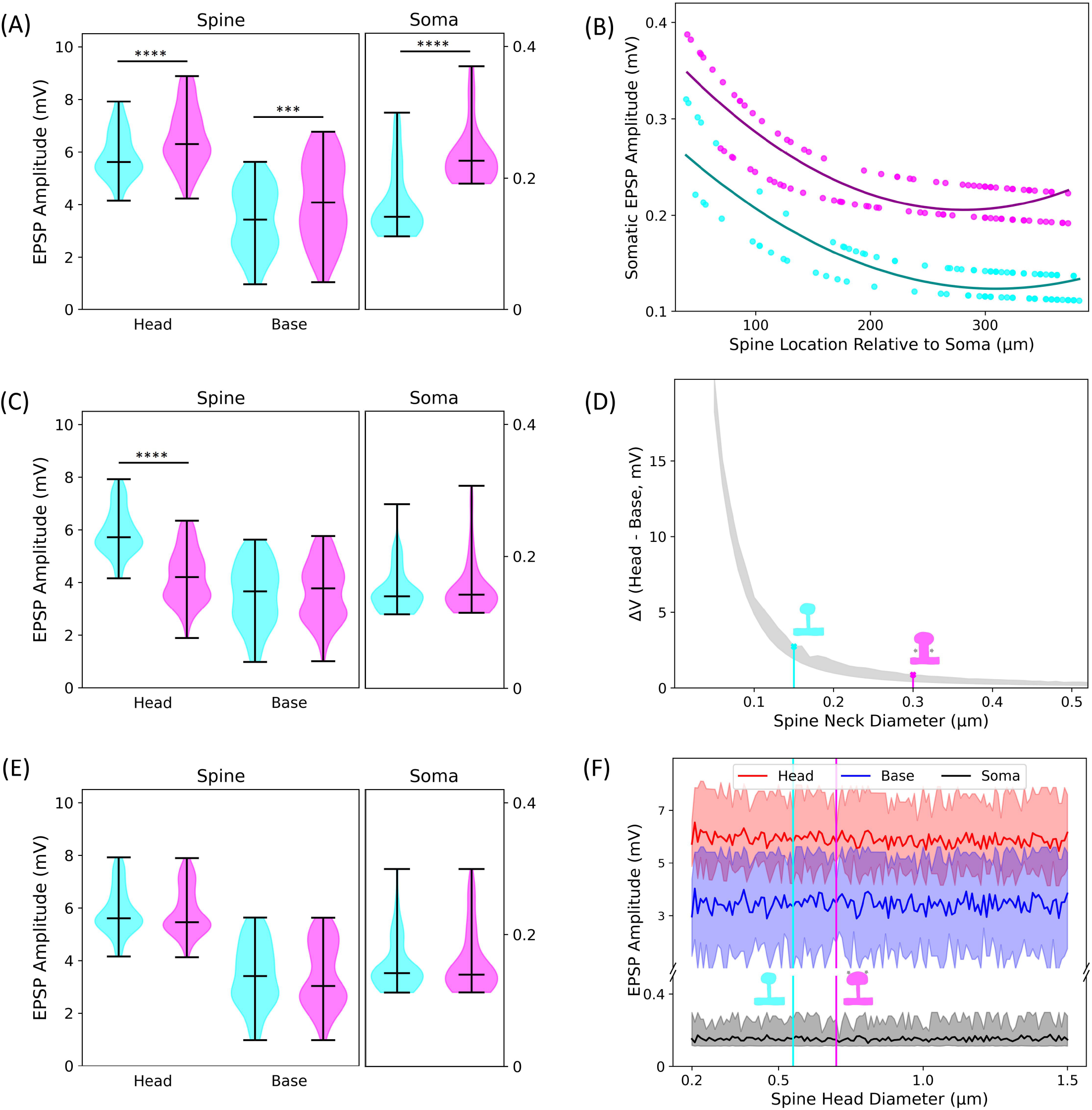
Differences in cellular responses associated with variations in dendritic spine morphology and density. (A, C, E) Violin plots of EPSP amplitudes recorded following single-spine activation in models differing in spine density, neck diameter, and head diameter, respectively. Data represent 100 simulations per group (N = 100); horizontal lines indicate the median. **** P < 0.0001, *** P < 0.001 (Mann– Whitney U test). (B) Relationship between the distance from the soma to the stimulated spine and the resulting somatic EPSP amplitude. Dotted lines represent individual simulation values; solid lines show linear regression fits for each model. (D) Voltage difference between the spine head and base as a function of spine neck diameter, based on 20 simulations per condition. (F) EPSP amplitude as a function of spine head diameter, with 20 simulations performed per condition.

We next examined how individual spine morphology affects signal propagation. Doubling the spine neck diameter in the altered model significantly reduced EPSP amplitude at the spine head but not at the base or soma (Figure 2C, Supplementary Figure 1D). Voltage differences between spine head and base declined rapidly with increasing neck diameter, becoming negligible above 0.4µm (Figure 2D), indicating that neck geometry critically regulates local electrical compartmentalization. In contrast, variations in spine head diameter, even across a 0.2–1.5µm range, had minimal impact on EPSP amplitude at any site (Figure 2E, Figure 3F, Supplementary Figure 1E). These findings highlight the dominant role of spine neck morphology in shaping dendritic voltage propagation, whereas spine head size has little effect on signal conduction.

**Figure 3.**
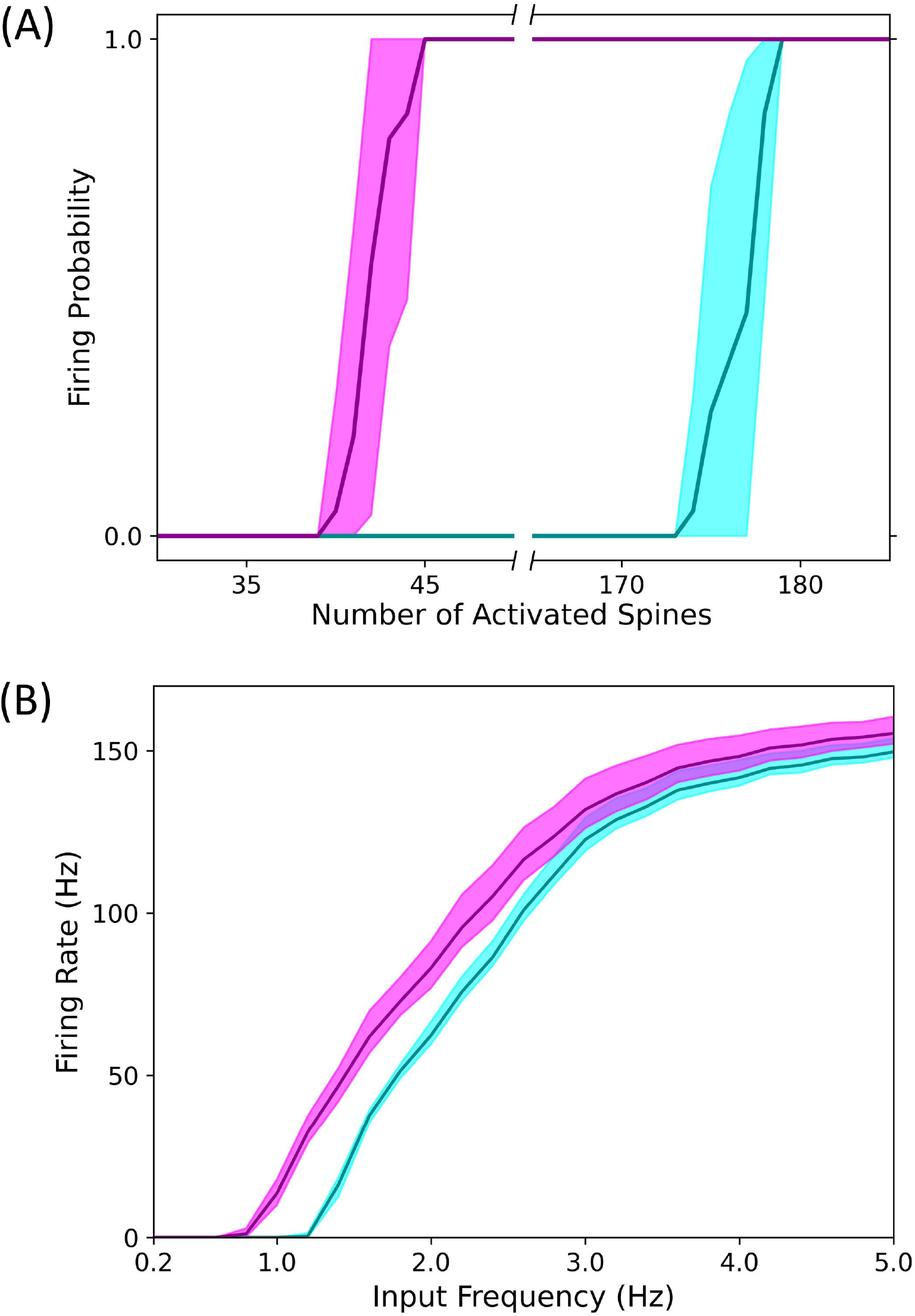
Comparison of cell-level excitability between models with different spine architectures. (A) Firing probability as a function of the number of simultaneously stimulated spines. Each data point represents the results of 20 simulations per spine count for each model. (B) Input–output firing rate curves in response to stochastic synaptic input delivered independently to explicit spines. Solid lines represent the mean firing rate. Each input rate was tested with 30 simulations per model.

### Morphological Alterations Promote Hyperexcitability

To assess how single-spine responses affect neuronal excitability, we measured the action potential threshold during synchronous spine activation. The baseline model required activation of 174 spines to trigger an action potential, whereas the altered model spiked with only 40 spines (Figure 3A). Considering the total number of explicit spines, this corresponded to 10.4% activation in the baseline model and 6.6% in the altered model. These findings demonstrate a markedly reduced spiking threshold in the altered model, indicating heightened excitability at the level of convergent synaptic input.

We next examined how these differences manifest under stochastic synaptic input. Independent Poisson-distributed synaptic events were delivered to explicit spines while systematically varying the relative contributions of apical and basal dendrites. Across the full tested range of input frequencies (0.2–5Hz), the altered model exhibited higher mean firing rates than the baseline model (Figure 3B). At higher input rates (≥3Hz) the difference remained modest, with the altered model firing at an average of 1.04 times the baseline rate, whereas under sparse input (<2Hz) the disparity became more pronounced, reaching 1.9-fold higher firing rates.

Together, these results demonstrate that the altered model exhibits heightened excitability and greater responsiveness to weak, distributed synaptic input.

## Discussion

An imbalance between E-I signaling has long been implicated in epileptogenesis^24,25^. Recently, volume EM has revealed ultrastructural changes in excitatory compartments of epileptic neurons, including hypertrophic spines, mitochondrial abnormalities, and loss of spine apparatuses^15^. Yet, direct measurement of dendritic spine signal transmission remains technically challenging due to their small size, heterogeneity, and complex biophysics^26,27^. To address these limitations, we used computational models to simulate pathological spine conditions observed in FCD (Figure 1). This approach enabled isolation of individual structural factors. The altered model showed weakened electrical compartmentalization at the spine neck and reduced spine density, which collectively decreased passive conductance channels and surface area, diminished signal attenuation, and increased depolarization spread toward the soma (Figure 2). These changes produced elevated neuronal excitability and lowered spiking thresholds under both controlled and stochastic inputs (Figure 3). Our models incorporated EM–derived spine morphologies rather than relying on idealized shapes, thereby capturing biologically realistic microstructural features and avoiding oversimplified representations. These results indicate that alterations in excitatory microstructure alone can elevate neuronal excitability, extending the traditional view of epilepsy pathophysiology and suggesting new targets beyond GABAergic modulation or Na+/Ca^2^+ channel blockade.

This study isolates the structural contributions to excitability, while other processes such as Ca^2+^ dynamics and intracellular signaling cascades^28–30^ were beyond scope. Consequently, features like loss of spine apparatuses^15^, which regulate Ca^2+^ buffering and compartmentalization^31,32^, were not represented due to their complexity^33^. The altered model also relied on population-averaged morphologies and excluded hypertrophic giant spines reported in epilepsy tissue^15^. These considerations suggest that our findings likely provide a conservative estimate of excitability, highlighting the robustness of the identified structural mechanisms.

Taken together, this study demonstrates that pathological modifications in dendritic spine structure, specifically reductions in density and neck compartmentalization can independently drive increased neuronal excitability spike output. These findings suggest a potential mechanism underlying epileptogenesis and highlight the importance of excitatory microstructure in shaping neuronal behavior. Expanding this work to the network of neurons, particularly in the context of the epileptic neurons contained in between control ones, will be essential for better understanding how microstructural alterations scale up to circuit level hyperexcitability and epileptic dynamics.

## Funding Statement

This research was supported by the KBRI Basic Research Program (25-BR-01-01, 25-BR-01-03) and the Brain Science Leading Convergence Technology Development Program (RS202300265524) of KBRI funded by the Ministry of Science and ICT.

## Conflict of Interest

None of the authors has any conflict of interest to disclose.

---

**Supplementary Figure 1.** Single-spine-evoked EPSPs recorded at three subcellular locations under distinct dendritic spine morphologies. (A) Schematic of recording sites: spine head (red), spine base (blue), and soma (black). EPSPs were recorded under five conditions: (B) baseline, (C) altered spine density only, (D) altered spine neck diameter only, (E) altered spine head diameter only, and (F) all spine architectural parameters altered simultaneously. For each condition (B–F), the top row shows raw EPSP traces from all three sites, and the bottom row shows an expanded view of the somatic EPSP. Light-colored traces represent individual trials (20 simulations per condition), and darker traces denote the trial-averaged responses. Horizontal cyan dashed lines indicate the peak EPSP amplitude in the baseline condition.

## Notes

### Competing Interest Statement

The authors have declared no competing interest.

