## Supplementary figures and images for "Microstructural Spine Alterations Increase Neuronal Excitability in Focal Cortical Dysplasia"

### Supplemental Figure 1

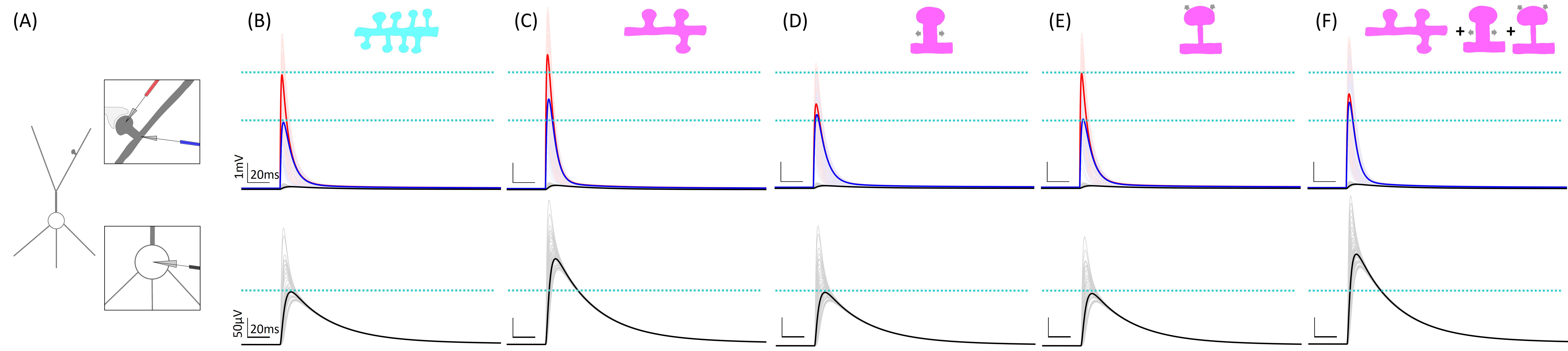
